# Optogenetic EB1 inactivation shortens metaphase spindles by disrupting cortical force-producing interactions with astral microtubules

**DOI:** 10.1101/2021.09.29.462463

**Authors:** Alessandro Dema, Jeffrey van Haren, Torsten Wittmann

## Abstract

Chromosome segregation is accomplished by the mitotic spindle, a bipolar micromachine built primarily from microtubules. Different microtubule populations contribute to spindle function: Kinetochore microtubules attach and transmit forces to chromosomes, antiparallel interpolar microtubules support spindle structure, and astral microtubules connect spindle poles to the cell cortex [1,2]. In mammalian cells, End Binding (EB) proteins associate with all growing microtubule plus ends throughout the cell cycle and serve as adaptors for a diverse group of +TIPs that control microtubule dynamics and interactions with other intracellular structures [3]. Because binding of many +TIPs to EB1 and thus microtubule-end association is switched off by mitotic phosphorylation [4–6] the mitotic function of EBs remains poorly understood. To analyze how EB1 and associated +TIPs on different spindle microtubule populations contribute to mitotic spindle dynamics, we use a light sensitive EB1 variant, π-EB1, that allows local, acute and reversible inactivation of +TIP association with growing microtubule ends in live cells [7]. We find that acute π-EB1 photoinactivation results in rapid and reversible metaphase spindle shortening and transient relaxation of tension across the central spindle. However, in contrast to interphase, π-EB1 photoinactivation does not inhibit microtubule growth in metaphase, but instead increases astral microtubule length and number. Yet, in the absence of EB1 activity astral microtubules fail to engage the cortical dynein/dynactin machinery and spindle poles move away from regions of π-EB1 photoinactivation. In conclusion, our optogenetic approach reveals mitotic EB1 functions that remain hidden in genetic experiments likely due to compensatory molecular systems regulating vertebrate spindle dynamics.

## RESULTS AND DISCUSSION

### π-EB1 photoinactivation reversibly shortens metaphase spindles

To test how acute inactivation of EB1-mediated +TIP recruitment to growing microtubule (MT) plus ends affects mitotic spindle organization and dynamics, we utilized our previously published H1299 cell line in which the endogenous EB1 and EB3 genes are knocked out by CRISPR/Cas9 genome editing and replaced by stable expression of blue light sensitive π-EB1 [7]. π-EB1 has its N- and C-terminal domains separated by a LOV2/Zdk1 module that in response to blue light results in rapid and reversible dissociation of the π-EB1 N-terminal MT- and C-terminal +TIP-binding activities. In cells in which the C-terminal half of π-EB1 is tagged with mCherry, mCherry-Zdk1-EB1C associates with growing plus ends of both spindle as well as astral MTs in the absence of blue light and rapidly dissociates from MT ends upon blue light exposure (Fig. 1A) similar to what we described in interphase cells [7].

**Figure 1.**
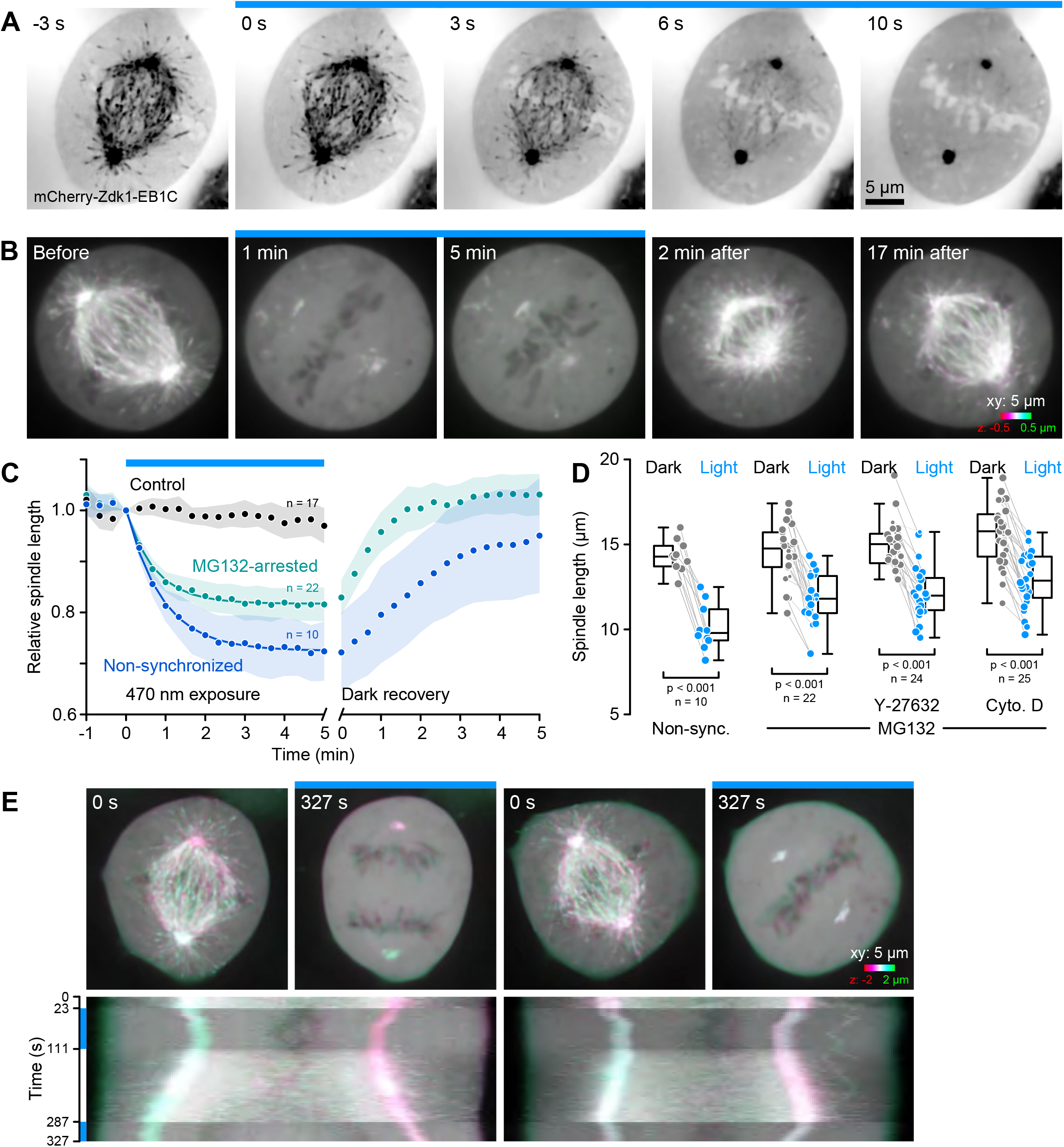
π-EB1 photoinactivation causes rapid and reversible metaphase spindle shortening. **(A)** Metaphase π-EB1 H1299 cell demonstrating blue light-mediated rapid dissociation of the mCherry-tagged π-EB1 C-terminal half from growing MT ends, but not from spindle poles. Single channel images in this and subsequent figures are shown with inverted contrast for improved visibility. **(B)** Time-lapse of a similar cell over a longer period demonstrating spindle shortening during blue light exposure and elongation after blue light exposure is terminated. Images are color-coded maximum intensity projections to compensate for spindle pole movements out of focus. **(C)** Relative spindle length over time in response to blue light. Shown is the average of the indicated number of cells for each condition. Shaded areas are 95% confidence intervals, and solid lines single exponential fits of the shortening phase. **(D)** Spindle length comparison before and during blue light exposure in the indicated conditions. In this and subsequent figures, box plots show median, first and third quartile, with whiskers extending to observations within 1.5 times the interquartile range. The size of the individual data points shows the goodness of the exponential fit and grey lines connect data points from the same cell. Statistical analysis by paired Student’s t-test. **(E)** Color-coded maximum intensity projections of the first and last frames and kymographs along the pole-to-pole axis of a time-lapse sequence of two cells from the same field of view. The cell on the left enters anaphase during the dark recovery phase. Note that anaphase elongation does not stop during a second blue light exposure. Blue bars in all panels indicate time phases with blue light stimulation.

Given that spindles are shorter downstream of EB1 RNA interference in *Drosophila* S2 cells [8] and that a fraction of mCherry-Zdk1-EB1C remains at the spindle poles during blue light exposure independent of MT-binding [9], we first analyzed spindle length in response to acute π-EB1 inactivation. Indeed, spindles rapidly shortened immediately following blue light illumination (Fig. 1B; Video 1). On average this shortening response was well described by a first order exponential decay with a shortening halflife of ~40 seconds (Fig. 1C), and using exponential fitting of individual spindle time-lapse sequences to calculate a new steady-state length during blue light exposure, we found that spindles shortened by ~30% (Fig. 1D). In contrast to this metaphase spindle shortening, π-EB1 did not block anaphase spindle elongation (Fig. 1E; Video 2). We therefore arrested and enriched cells in metaphase with the proteasome inhibitor MG132 that prevents sister chromatid separation [10]. π-EB1 photoinactivation induced spindle shortening in these metaphase-arrested spindles with similar kinetics, although metaphase-arrested spindles remained slightly but significantly longer during blue light compared with spindles in non-arrested cells (Fig. 1C, D). Nevertheless, because the response was more consistent, we used the MG132 block in most subsequent experiments. Spindle shortening was reversible, and with or without MG132, spindles returned to their original length after blue light exposure (Fig. 1C, E). In addition, spindles in control cells not expressing π-EB1 did not respond to blue light demonstrating that spindle shortening was not due to phototoxicity (Fig. 1C; Fig. S2), but instead is a specific effect of acute EB1 inhibition. To test if different populations of spindle MTs contribute to spindle length, we inactivated π-EB1 by localized blue light exposure at either the cell cortex or the metaphase plate. In both types of experiments, spindles shortened similarly, but to a smaller extent compared with whole cell π-EB1 photoinactivation (Fig. S1A). This suggested that both astral and spindle MTs contribute to spindle length maintenance although diffusion of photoinactivated EB1 to other parts of the cell cannot be excluded. Neither inhibition of actomyosin contractility with the Rho-kinase inhibitor Y27632 or a low dose of cytochalasin D that inhibits actin polymerization dynamics in mitosis [11] altered the spindle shortening response (Fig. 1D) implying that +TIP-mediated spindle length maintenance is predominantly MT driven. Interestingly, spindles in EB1/EB3 −/− cells were not shorter than in control cells (Fig. S1C) indicating that genetic loss of EB1 can be compensated and highlights the necessity of acute perturbation to reveal mitotic EB1 functions.

### EB1 activity is required for astral microtubule length control

Inhibition of MT dynamics with paclitaxel results in rapid spindle shortening possibly by inhibition of kinetochore (KT)-associated MT assembly while maintaining MT depolymerization near the spindle poles [12–14]. Remarkably, in H1299 cells the kinetics and extent of spindle shortening either with paclitaxel (Fig. S1B) or by π-EB1 photoinactivation were remarkably similar. In interphase, π-EB1 photoinactivation reduces the MT growth rate and increases catastrophe frequency [7]. Thus, to test if inhibition of MT growth causes the observed spindle shortening phenotype, we asked how π-EB1 photoinactivation affected MT dynamics in mitotic cells. Computational tracking of the fluorescently tagged N-terminus of π-EB1, which remains on MT ends during blue light exposure, revealed that the MT growth rate in metaphase was substantially lower than in interphase [7]. In addition, and in contrast to interphase, the metaphase MT growth rate was only minimally reduced by π-EB1 photoinactivation (dark: 15.1 +/− 2.3 μm/min; blue light: 14.3 +/− 2.4 μm/min; Fig. 2A; Video 3), and not statistically significantly different at the 99% significance level. This indicates that EB1-dependent mechanisms that sustain fast growth of cell body MTs in interphase are absent in mitosis, but also shows that a decrease in MT growth cannot explain the observed spindle shortening phenotype.

**Figure 2.**
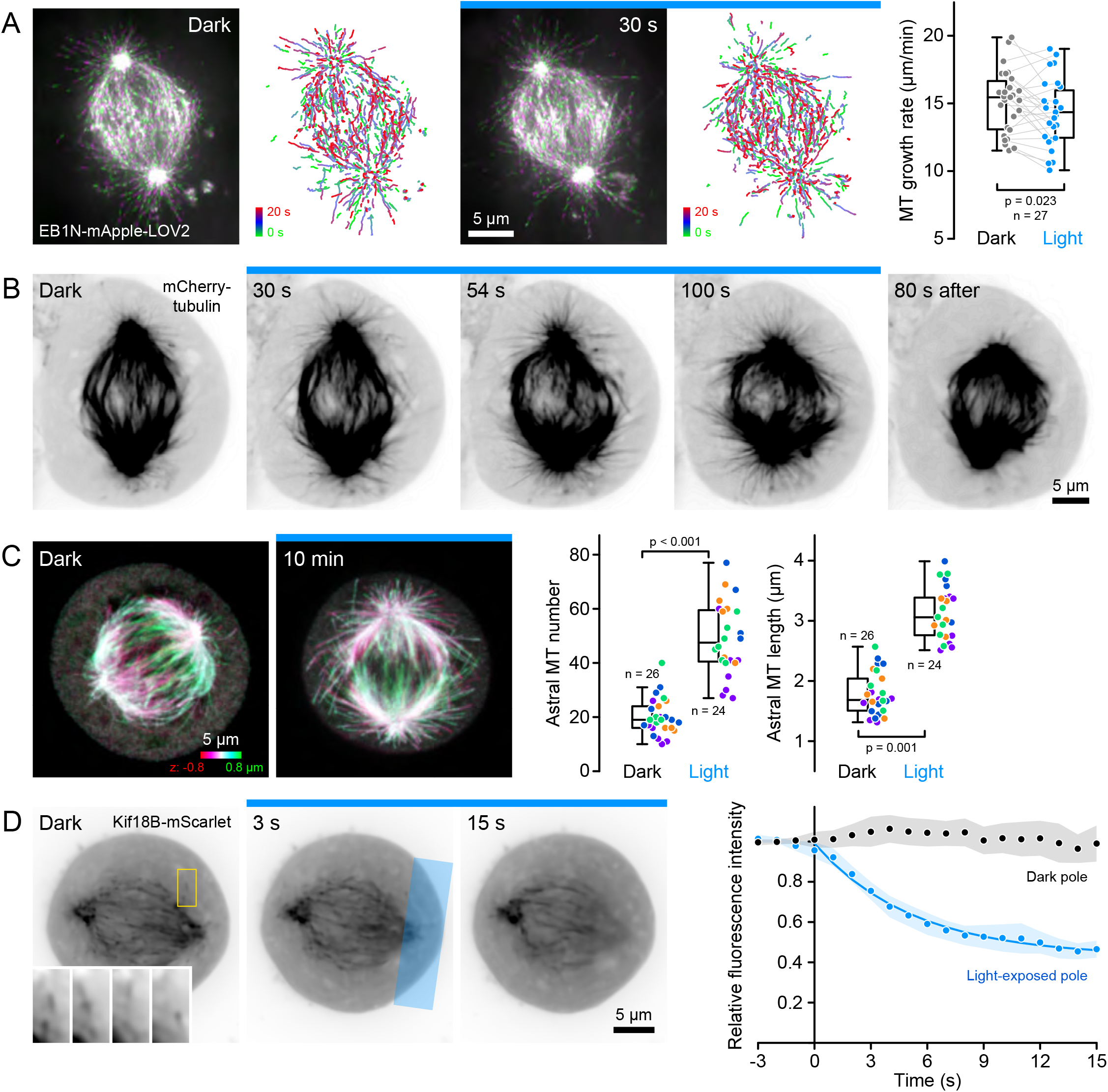
π-EB1 photoinactivation increases astral MT length. **(A)** EB1N-mApple-LOV2-labelled MT plus ends before and during blue light exposure in MG132-arrested metaphase spindles. Maximum intensity projections in alternating colors over 20 s at 3-s intervals and MT growth tracks over the same time window illustrate MT growth dynamics before and during blue light exposure. The comparison of the median MT growth rate per cell before and during blue light exposure is shown on the right. Grey lines connect data from the same cell. Statistical analysis by paired Student’s t-test. **(B)** Time-lapse sequence of a π-EB1 H1299 cell expressing mCherry-tagged α-tubulin. Note the increase in astral MT number and length yielding a ‘hairy’ spindle appearance during blue light exposure, and that these ectopic MTs rapidly disappear when blue light is switched off. **(C)** Color-coded maximum intensity projections of MTs in fixed metaphase spindles in the dark or after 10 min blue light exposure. Box plots show the comparison of astral MT length and number. Data from 4 independent experiments are indicated by different colors. Statistical analysis by unpaired Student’s t-test of the experimental mean. **(D)** π-EB1 H1299 cell expressing mScarlet-I-tagged Kif18B in the dark and during local blue light exposure of the right spindle pole (indicated by the blue box in the middle panel). Inset in the left panel shows KIF18B-mScarlet localization to growing astral MT ends at 2 s intervals. The graph shows the relative KIF18B-mScarlet fluorescence intensity over time comparing the non-illuminated and the blue light-exposed pole in n = 19 cells with the shaded areas indicating 95% confidence intervals and the solid blue line an exponential fit. Single channel images in B and D are shown with inverted contrast, and blue bars in all panels indicate time phases with blue light stimulation.

To test how EB1 supports spindle MT organization independent of growth rate changes, we directly observed mCherry-tubulin dynamics in π-EB1 expressing cells. To our surprise and quite opposite to MT growth inhibition, blue light exposure quickly induced an increase of MTs originating from the spindle poles resulting in an overall ‘hairy’ appearance even before spindle shortening became apparent (Fig. 2B; Video 4). This effect was reversible and MTs extending into the surrounding cytoplasm quicky disappeared once blue light exposure was terminated. Because the high background of soluble mCherry-tubulin in mitosis makes it difficult to resolve astral MT ends in live cells, we quantified this effect by comparing spindle MTs in cells fixed in the dark and during blue light exposure. In these experiments, in which a slightly lower dose of blue light was delivered to the whole coverslip using a custom-designed LED ring [15] spindle shortening remained fully reversible after 20 mins of blue light exposure (Fig. S2). Remarkably, both the number and length of astral MTs increased dramatically during blue light exposure compared with cells that were kept in the dark with many astral MTs extending to the cell cortex (Fig. 2C). While we cannot exclude that unbundling of antiparallel central spindle MTs contributes to this hairy spindle phenotype [16], the observed increase in astral MT length is very similar to the RNA interference phenotype of KIF18B [17,18], a MT-destabilizing kinesin implicated in astral MT length control in metaphase [18,19]. KIF18B binds to EB1 through a SxIP motif [19], and indeed, in H1299 π-EB1 cells, KIF18B-mScarlet associated with growing astral MT ends in the dark but was rapidly lost from spindle MTs during blue light exposure (t1/2 = ~4 s, Fig. 2D). Thus, although *in vitro* KIF18B can bind MTs independent of EB1 [20], our results demonstrate that KIF18B-mediated astral MT pruning in metaphase requires EB1.

### EB1 is required to engage cortical pulling forces in metaphase

Spindle position and orientation are controlled by pulling forces on astral MTs by dynein/dynactin localized to the cell cortex [21]. Transition from end-on MT interactions with this cortical force-generating machinery in metaphase to lateral dynein/dynactin-binding along the side of astral MTs in anaphase is thought to increase dynein-mediated forces partially driving anaphase spindle elongation. Indeed, experimentally producing lateral MT attachment in metaphase increases spindle length [22], the opposite of the spindle shortening we observe when inhibiting π-EB1. Consistent with the hypothesis that π-EB1 photoinactivation disrupts astral MT interactions with cortical dynein/dynactin in metaphase, blue light exposure of only one spindle pole in H1299 π-EB1 cells resulted in asymmetric spindle shortening in which the blue light illuminated pole moved away from the cell cortex while the non-illuminated pole remained unaffected (Fig. 3A; Video 5). In addition, while spindle orientation fluctuated randomly in the dark, asymmetric π-EB1 photoinactivation on opposite sides of the pole-to-pole axis at the cell cortex near each spindle pole resulted in a spindle rotation bias away from the blue light exposed regions (Fig. 3B). Although spindle orientation continued to fluctuate randomly after blue light exposure was terminated, on average spindles remained in their new positions. Because in these experiments, spindle rotation stopped once the spindle poles had moved away from the blue light exposed regions, to test if spindles could be rotated further, we then slowly moved wedge-shaped π-EB1 photoinactivation regions on a circular path around the cell cortex by 1° every 40 seconds. This experiment was technically difficult and only approximately a third of the cells responded clearly to the rotating blue light pattern. However, in the cells that did, the spindle rotation angle followed the rotation ahead of the blue light pattern closely providing further evidence that off-axis π-EB1 photoinactivation generates an imbalance of cortical pulling forces (Fig. 3C; Video 5). Faster movement of the photoinactivation regions did not result in faster spindle rotation. Instead, the blue light pattern passed by the spindle poles indicating a limiting rate at which spindles can rotate. Together with our data that π-EB1 photoinactivation does not cause anaphase spindle shortening (Fig. 1E), this shows that EB1 is required to engage the force-generating dynein/dynactin machinery in metaphase, but not in anaphase.

**Figure 3.**
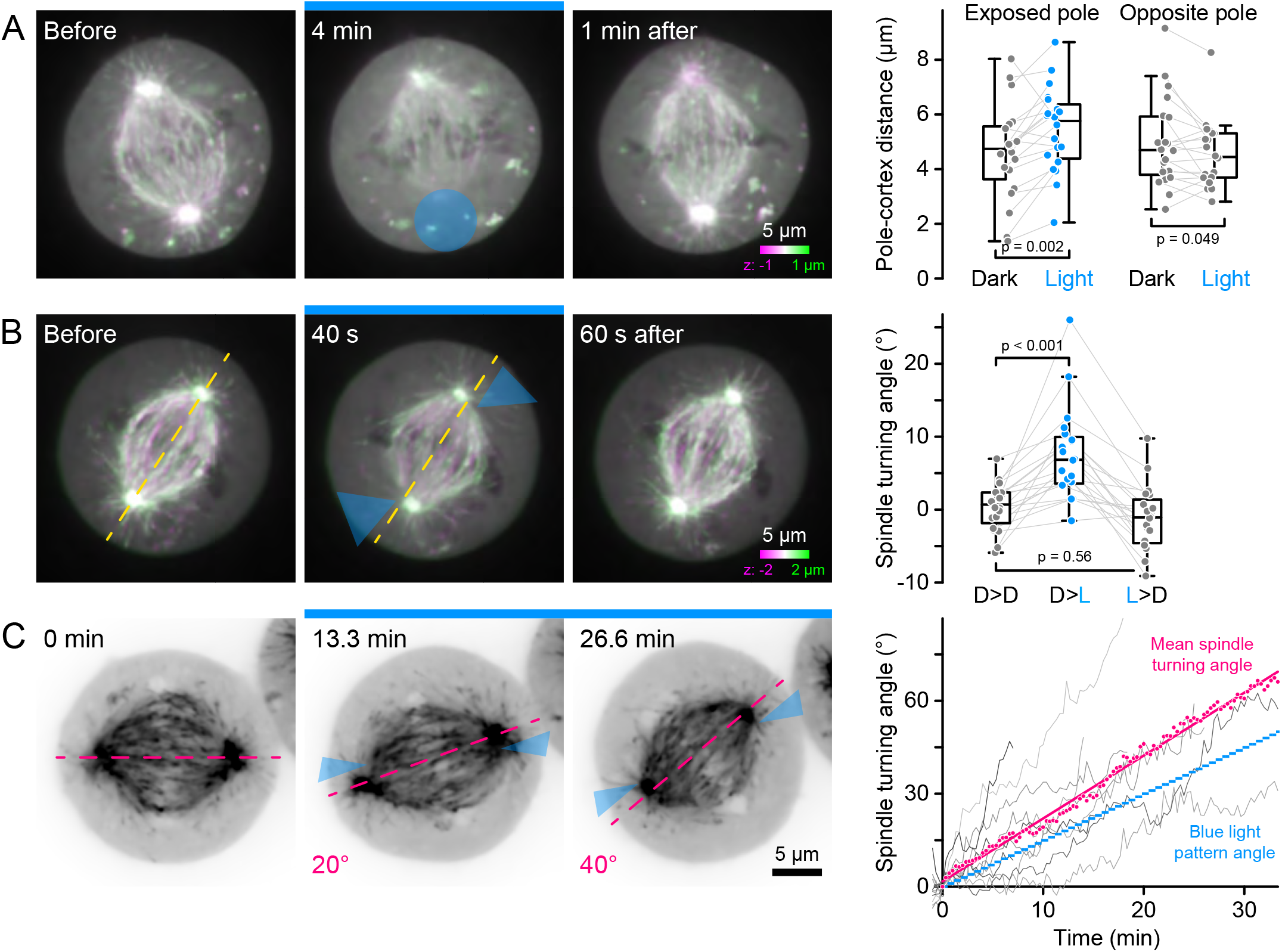
Spindle rotation by local π-EB1 photoinactivation. **(A)** Metaphase π-EB1 H1299 cell in which only the lower spindle pole was exposed to blue light. Box plots show the nearest distance of both the light-exposed and the opposite spindle pole from the cortex in the dark and 2 min during localized blue light exposure (n = 20 cells). Statistical analysis by paired Student’s t-test. **(B)** Metaphase π-EB1 H1299 cell with off-axis blue light exposure near both spindle poles. Box plots show the degree of spindle rotation over 1 min intervals in the dark (D>D), when blue light is switched on (D>L), and when blue light is switched off (L>D) with positive angles defined as rotation away from the blue light pattern (n = 20 cells). Statistical analysis by one-way ANOVA and Tukey-Kramer HSD test. **(C)** Metaphase π-EB1 H1299 cell with off-axis blue light pattern rotation by 1° every 40 s. The magenta dashed line shows the expected rotation at the indicated time points. In the graph, grey lines are the spindle axis rotation of individual cells that responded to the rotation. Magenta symbols are the average rotation of these n = 11 cells with the solid line showing a linear fit. The blue stepped line shows the rotation of the blue light pattern. Images in A and B are color-coded maximum intensity projections, and in C are shown with inverted contrast. All cells were arrested in metaphase with MG132. and localized blue light exposure is indicated by blue transparent regions.

There are two possibilities how π-EB1 photoinactivation might interfere with cortical pulling forces. More and longer astral MTs may physically resist spindle pole movement toward the cortex, or EB1 may also be required for dynein/dynactin-mediated cortical MT capture. While we cannot distinguish between these mechanisms and both may be at play, it is important to note that the dynactin subunit 1 (DCTN1 or p150Glued) can itself be a +TIP, and exists in alternatively spliced variants [23,24]. While the longer neuronal DCTN1 isoform binds directly to MTs, in non-neuronal cells a 20 amino acid section of the basic MT-binding domain is removed reconstituting a cryptic SKLP motif required for binding to EB1 (Fig. S3A). Indeed, both in interphase and mitotic cells, only plus-end-tracking and spindle localization of the shorter non-neuronal DCTN1 isoform was sensitive to π-EB1 photoinactivation while association of the neuronal isoform along MTs was unaffected (Fig. S3B). Although we were unable to visualize DCTN1 directly on astral MT ends, this suggests that in addition to recruiting KIF18B EB1 contributes to productively link dynein/dynactin to growing astral MT ends.

### π-EB1 photoinactivation transiently relaxes inter-kinetochore tension

In metaphase, the two KTs of correctly attached chromatid pairs are pulled toward opposite spindle poles and the stretch between sister KTs can serve as relative readout of tension across the central spindle [6,25]. Thus, to test how loss of cortical pulling on astral MTs is transmitted through the spindle, we asked how the distance between sister KTs changes in response to π-EB1 photoinactivation. Even though this inter-KT distance fluctuates substantially over time, KT pairs in π-EB1 cells frequently appeared to shorten and twist away from the spindle axis during blue light exposure (Fig 4A). Because it was difficult to accurately follow individual sister KT pairs over time, we instead measured the length of as many KT pairs as could be identified in single optical sections at specific time points during blue light exposure. At 3 min, the average cellular sister KT-KT distance was significantly reduced by ~13% (dark: 0.83 +/− 0.11 μm; blue light: 0.72 +/− 0.11 μm; Fig. 4B) indicating that spindles acutely relax in response to disengaging cortical pulling forces. However, this relaxation was transient and normal KT-KT distance was quickly restored in the absence of EB1 activity demonstrating an adaptive response of other EB1-independent force generating mechanisms. In addition, KT pairs did not shorten in response to localized π-EB1 photoinactivation of only a small KT population on one side of the spindle (Fig. 4C). Together, this implies that EB1 is not required for metaphase KT attachment even though it is localized to growing KT MT bundle ends [26]. Nevertheless, π-EB1 photoinactivation did impact progression through mitosis. Although π-EB1 H1299 cells entered mitosis normally during prolonged π-EB1 photoinactivation and most cells eventually divided, metaphase spindles remained slow and the duration from nuclear envelope breakdown (NEB) to completion of metaphase was ~3 times longer compared with cells that were kept in the dark (Fig. 4D, E).

**Figure 4.**
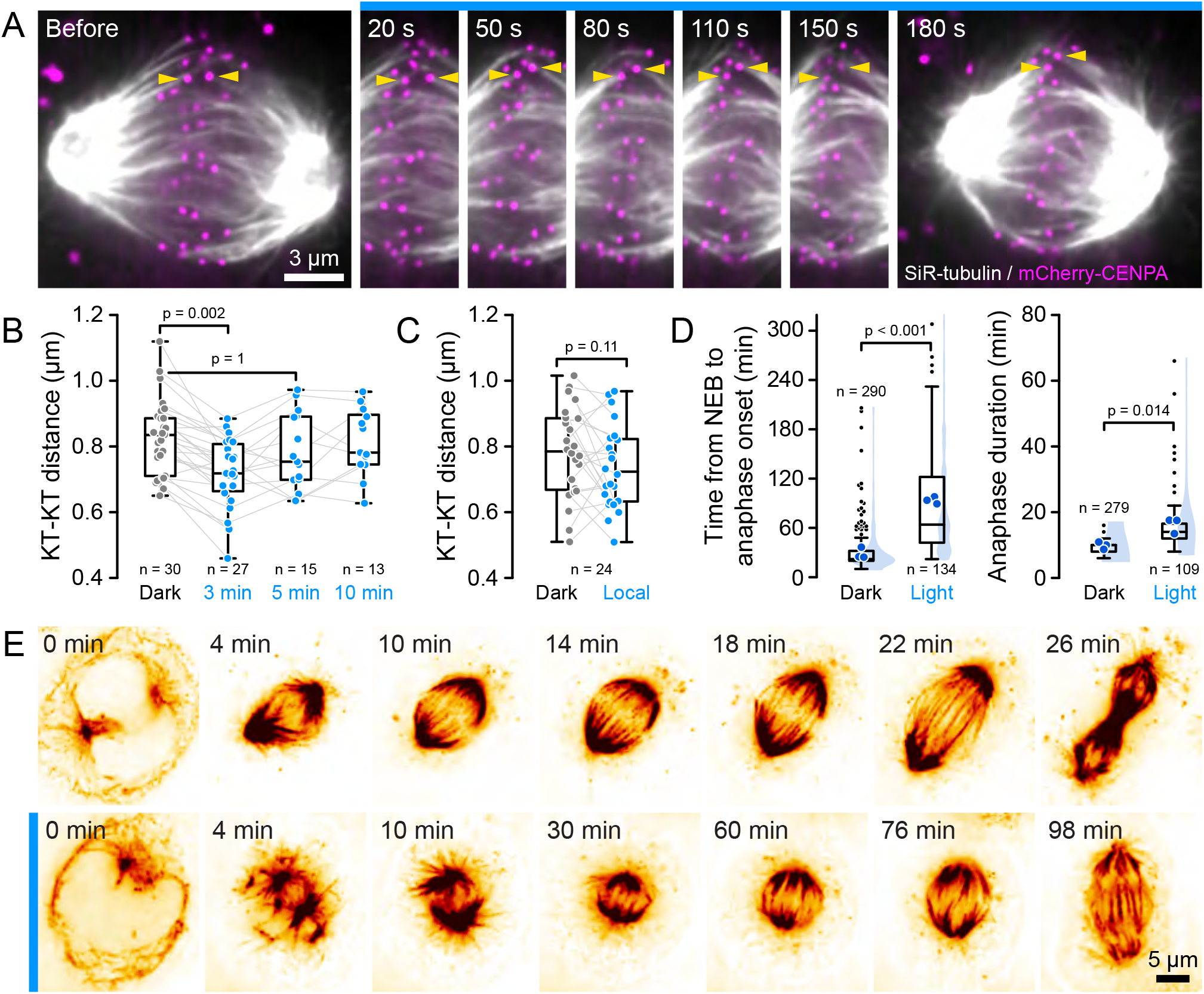
π-EB1 photoinactivation induces transient relaxation of inter-KT tension. **(A)** Time-lapse sequence of a metaphase π-EB1 H1299 cell expressing mCherry-CENPA to label KTs. MTs are labeled with SiR-tubulin. Yellow arrowheads highlight a KT pair that is easy to follow and visibly shortens during blue light exposure. **(B)** Distance between sister KTs in π-EB1 H1299 metaphase spindles in the dark and at indicated times during blue light exposure. Grey lines connect data points from the same cells but note that not all cells were followed through all time points. Statistical analysis by one-way ANOVA and Tukey-Kramer HSD test indicating significant KT-KT distance shortening only at 3 min. **(C)** Comparison of distance between sister KTs in the dark and after 3 min of local blue light exposure of only a few KT pairs. Statistical analysis by paired Student’s t-test. Grey lines connect data points from the same cells. **(D)** Timing of progression through mitosis in π-EB1 H1299 in the dark or exposed to blue light. The blue shaded areas show the kernel density distribution of all observed cells from 3 independent experiments. Statistical analysis by unpaired Student’s t-test of the experimental mean (dark blue symbols). **(E)** Examples of π-EB1 H1299 cell progression through mitosis in the dark and during blue light exposure. MTs were labeled with SPY555-tubulin. Blue bars in all panels indicate time phases with blue light stimulation.

Because accurate chromosome segregation is fundamental to life, redundant systems ensure mitotic spindle fidelity, consistent with the reported mild mitotic phenotypes of genetic EB1 removal in *Drosophila* [8,27] and the absence of dramatic spindle defects in vertebrate EB1/3 −/− cells (Fig. S1C) [28]. In contrast, by forestalling compensatory mechanisms, acute optogenetic π-EB1 inactivation reveals EB1 functions that remain hidden in genetic experiments. However, it should be noted that the π-EB1 dissociation products may have additional dominant effects and therefore more dramatic consequences than EB1 removal alone. For example, the EB1 C-terminal half could interfere with cortical MT capture, although mCherry-Zdk1-EB1C does not localize to the cortex in blue light. Notably, although MT growth has been linked to spindle length [29,30], π-EB1 photoinactivation-mediated spindle shortening was not accompanied by a MT growth rate decrease. Instead, we think that acute EB1 inhibition causes an imbalance of forces acting on the spindle poles that actively drives spindle shortening. To our knowledge, such fast and reversible spindle length change has not previously been reported. Spindles do not shorten during optogenetic inactivation of the MT-bundling protein PRC1 [31]. Consistent with pulling on astral MTs, spindles move toward optogenetically recruited NuMa [32], but also in these experiments spindle length does not change indicating that spindle tension persists. Spindles remained short in π-EB1 H1299 cells in blue light and only returned to their original length when EB1 activity is reinstated, in agreement with force-independent slower developmental scaling mechanisms that act through controlling component or MT nucleation site availability [30,33,34]. In conclusion, our results highlight the importance of EB1-mediated MT end interactions in regulating spindle length and position, which in addition to ensuring accurate chromosome segregation [35] have important roles in asymmetric developmental cell divisions [36].

## AUTHOR CONTRIBUTIONS

Conceptualization, A.D., J.v.H. and T.W.; Methodology, A.D., J.v.H. and T.W.; Software, T.W.; Investigation, A.D. and J.v.H.; Writing – Original Draft, A.D. and T.W.; Writing – Review & Editing, A.D., J.v.H., and T.W.; Funding Acquisition, T.W.; Supervision, T.W.

## ACKNOWLEDGEMENTS

We thank Marvin Tanenbaum and Michael Davidson for plasmids and the Dumont lab and all members of the HSW-6 community for productive discussions. This work was supported by National Institutes of Health grants R21 CA224194, R01 NS107480, S10 RR026758 and S10 OD028611 to T.W.

## METHODS

### Cell lines, culture and live cell treatments

Parental NCI-H1299 human non-small cell lung carcinoma cells (ATCC Cat# CRL-5803, RRID:CVCL_0060, male), or π-EB1 H1299 cells were cultured in RPMI 1640 supplemented with 10% FBS and non-essential amino-acids. Production of π-EB1 H1299 cells is described elsewhere [7,37]. H1299 cell lines were cultured at 37°C, 5% CO_2_ in a humidified tissue culture incubator and regularly tested for mycoplasma contamination (IDEXX BioResearch). H1299 cells and sublines were previously authenticated by STR profiling [7]. For microscopy, cells were plated in glass-bottom dishes (Mattek).

Cells were arrested in metaphase with 10 μM MG132 (Sigma Aldrich) shortly before imaging and were used for a maximum of 4 hours after MG132 addition. 10 μM Y27639 (Tocris) was added 2 hours before imaging to inhibit ROCK. 100 nM Cytochalasin D (Thermo Fisher) was added 1 hour before imaging to inhibit f-actin polymerization dynamics. 5 μM paclitaxel (Thermo Fisher) was added during imaging. Spy555-tubulin (Cytoskeleton. Inc) was added to the medium 30 minutes before cell imaging at a 1:2000 dilution from a stock prepared according to the manufacturer’s directions. SiR-tubulin (Cytoskeleton. Inc) was employed in a similar fashion at a 300 nM final concentration. Transfection was performed with Lipofectamine 3000 (Thermo Fisher) according to manufacturer’s protocol.

### Molecular cloning

The KIF18B coding sequence was amplified by PCR from 24xMoonTag-kif18b-24xPP7 (Addgene plasmid #128604 [38] was a gift from Marvin Tanenbaum) and cloned into KpnI and BamHI sites of pCMV-CKAP5-mScarlet-I [7] by Gibson Assembly thereby replacing the CKAP5 ORF with KIF18B.

pEB1N-mApple-LZ-LOV2 was cloned by inserting the mApple and LZ-LOV2 coding sequences into the XhoI and BamHI sites of EB1N-mCherry-LZ-LOV2 [7] by Gibson Assembly. mApple and LZ-LOV2 coding sequences were amplified using primers detailed in the key resource table.

mCherry-Dynactin-C-18, representing the neuronal DCTN1 isoform, was obtained from the UCSF Michael Davidson plasmid collection. The alternatively spliced ubiquitous DCTN1 isoform was generated by PCR amplification of the dynactin coding sequence upstream and downstream of the SKLP motif sequence using mCherry-Dynactin-C18 as a template. Resulting PCR amplicons were joined by overlap extension PCR using the two outermost primers. The final PCR product was digested with XhoI and EcoRI and ligated into the XhoI/EcoRI sites of plasmid pBio-mCherry-C1.

All constructs were verified by sequencing and primer sequences are included in the key resource table.

### Microscopy and π-EB1 photoinactivation

In most experiments, fluorescent protein dynamics were imaged by Yokogawa CSU-X1 spinning disk confocal microscopy on a microscope system essentially as described [39], but upgraded with a more sensitive Prime BSI sCMOS camera (Teledyne Photometrics) with a 60x 1.49 NA oil immersion lens (Nikon). On this microscope, a Polygon 1000 digital micromirror device (Mightex) equipped with a 470 nm LED is coupled into an auxiliary eyepiece camera port on a Ti-E inverted microscope stand (Nikon) [15]. Using an 80/20 beamsplitter in the emission lightpath this allows simultaneous confocal imaging and π-EB1 photoinactivation. Blue light exposure regions were drawn on the fly with the PolyScan2 software (Mightex) using a still acquisition as template, except to induce prolonged spindle rotation. In the spindle rotation experiment, a sequence of small triangular regions on opposite sides of the spindle pole axis was pre-programmed in PolyScan2 and rotated around the cell center by 1° every 40 s. For fast time-lapse imaging with acquisition frequencies at or above 1 Hz, blue light exposure was directly triggered by the camera with a 40 ms delay after camera exposure and 10 ms blue light pulses. In experiments with slower time lapse acquisition, the Polygon was triggered with an Arduino microcontroller at 1-2 Hz.

Imaging of EB1N-mApple-LOV2 to measure MT growth rates was performed on a newer Yokogawa CSU-W1/SoRa spinning disk confocal system in SoRa mode on a Ti2 inverted microscope stand (Nikon), and images acquired with an ORCA Fusion BT sCMOS camera (Hamamatsu) with 2×2 binning to reduce photobleaching. These experiments had three phases (20 s image acquisition without blue light exposure, 30 s blue light exposure without image acquisition, and finally 20 s image acquisition with blue light) with images acquired every 0.5 s to capture equilibrium MT dynamics before and during blue light exposure. This system was similarly equipped with a Polygon 1000 (Mightex) through an auxiliary filter turret and LAPP illuminator (Nikon) and integrated control of imaging and photoactivation was through NIS Elements v5.3 software (Nikon).

Lastly, bulk blue light exposure of cells in glass-bottom 12-well plates (Cellvis) for long-term multi-point imaging of cell cycle progression and for analyzing spindle structure in fixed cells was done with a custom-built Arduino-controlled 470 nm LED cube [15]. In all cases, the minimal blue light dose to achieve π-EB1 photoinactivation was determined by observing dissociation of fluorescently labeled π-EB1 C-terminal constructs from growing MT ends [7].

### Immunofluorescence

Microtubule staining was performed as described [40]. In brief, cells grown on coverslips were washed in PBS, fixed in 0.25% glutaraldehyde in 80 mM K-PIPES, 1 mM EGTA, 1 mM MgCl_2_, then quenched three times in 0.1% NaBH_4_ in PBS. Fixed cells were permeabilized with 0.3% Triton X-100 in PBS, then blocked in 1% BSA in PBS, and stained with rat anti α-tubulin (Biorad, MCA77G) diluted 1:750 in 0.1% Triton X-100, 1% BSA in PBS.

### Image analysis

Spindle length and rotation measurements as well as astral MT number and length quantifications were performed manually in Fiji [41]. Spindle length was determined by measuring the pole-to-pole distance in optical sections and equilibrium spindle length during blue light exposure was determined by fitting the spindle length time-lapse data for each cell with a single exponential decay function using the Curve Fitting toolbox in MATLAB (Mathworks, Inc.).

MT growth rates were measured with u-track essentially as described [7] with the exception that Poisson-distributed shot noise was removed prior to tracking using the NIS Elements v5.3 Denoise.ai neural net (Nikon). In side-by-side comparisons, this resulted in substantial improvement of MT growth track quality. Frame-to-frame MT growth rates were extracted from the tracking results using a custom MATLAB script and the median of all these instantaneous velocities per cell was calculated to minimize the impact of outlier tracking errors.

Sub-resolution distances between sister KTs [6] were measured with a custom MATLAB script, in which two Gaussian functions are fitted along an intensity profile across interactively selected KT pairs in single optical sections.

Images in figures and videos were processed with NIS Elements v5.3 Denoise.ai (Nikon) to reduce shot noise. No non-linear contrast adjustments were made, and images are shown in pseudo-color or contrast-inverted as indicated in the figure legends.

### QUANTIFICATION AND STATISTICAL ANALYSIS

Details of statistical analysis including the type of test, p-values and numbers of biological replicates are provided within the relevant figures and figure legends. All statistical analysis was done in MATLAB (Mathworks, Inc.)., and graphs were produced in MATLAB and in Excel (Microsoft). In all figures, box plots show median, first and third quartile, with whiskers extending to observations within 1.5 times the interquartile range. No randomization, stratification or sample size estimation strategies have been employed. Unless indicated otherwise in the text, all data points from a given experiment were analyzed, and a p < 0.01 is considered statistically significant. For every experiment reported, at least 3 biological replicates with similar results were performed.

