## Supplemental Material for "Optogenetic EB1 inactivation shortens metaphase spindles by disrupting cortical force-producing interactions with astral microtubules"

**SUPPLEMENTARY FIGURE LEGENDS**

**Figure S1.** **(A)** Spindle length comparison in π-EB1 H1299 cells before and during localized blue light exposure either on the metaphase plate or the cell cortex. Transparent blue areas indicate the light-exposed regions. **(B)** Rapid spindle shortening in response to 5 µM paclitaxel treatment in H1299 cells. All experiments were in MG132 metaphase-arrested cells and images are shown with inverted contrast. The size of the data points shows the goodness of the exponential fit. Grey lines connect data points from the same cell. Statistical analysis by paired Student's t-test in A and B. **(C)** Comparison of spindle length and distance between sister KTs in control H1299 cells, H1299 cells in which both EB1 and EB3 were deleted by CRISPR/Cas9 genome editing and these EB1/3 -/- cells expressing π-EB1. Statistical analysis by one-way ANOVA and Tukey-Kramer HSD test indicating no significant difference between these conditions.

**Figure S2.** Reversible spindle shortening in MG132-arrested π-EB1 H1299 cells exposed to blue light using a custom-designed 470 nm LED array. Shown are images before, immediately after 20 min of blue light exposure, and after 30 min recovery in the dark. Quantification shows the same time points in π-EB1 H1299 cells and in control H1299 cells expressing EB3-mCherry. Grey lines connect data points from the same cell. Statistical analysis by one-way ANOVA and Tukey-Kramer HSD test.

**Figure S3.** **(A)** Alignment of the MT-binding domain of neuronal and ubiquitous isoforms of DCTN1. Alternative splicing in the ubiquitously expressed DCTN1 isoform generates an EB1-binding SKLP motif highlighted in orange. **(B)** Change in DCTN1 localization in π-EB1 H1299 cells in response to blue light. Only the ubiquitous SKLP-motif DCTN1 isoform dissociates from spindle MTs in response to π-EB1 photoinactivation while spindle localization of neuronal DCTN1 was not affected. Although we were not able to visualize MT plus end localization of ubiquitous DCTN1 in mitosis, EB1-dependent plus-end-tracking was evident in interphase cells.

**SUPPLEMENTARY VIDEO LEGENDS**

**Video 1.** Time-lapse of a π-EB1 H1299 cell demonstrating spindle shortening during blue light exposure and elongation after blue light exposure is terminated. Images are color-coded maximum intensity projections to compensate for spindle pole movements out of focus. Related to Fig, 1B. Scale bar: 5 µm.

**Video 2.** Color-coded maximum intensity projections of two π-EB1 H1299 cells in the same field of view exposed to blue light as indicated. The cell on the left enters anaphase during the dark recovery phase. Note that anaphase elongation does not stop during a second blue light exposure. Related to Fig. 1E. Scale bar: 5 µm.

**Video 3.** EB1N-mApple-LOV2-labelled MT plus ends in an MG132 metaphase-arrested π-EB1 H1299 cell before and during blue light exposure. Note MT ends bumping into the cell cortex and failing to undergo catastrophes during blue light stimulation. Related to Fig. 2A. Scale bar: 5 µm.

**Video 4.** π-EB1 H1299 cell expressing mCherry-tagged α-tubulin exposed to blue light as indicated. Note the increase in astral MT number and length during blue light exposure, and that these ectopic MTs rapidly disappear when blue light is switched off. Related to Fig. 2B. Scale bar: 5 µm.

**Video 5. (A)** Metaphase π-EB1 H1299 cell in which only the lower spindle pole was exposed to blue light as indicated and moves away from the cortex during blue light exposure. **(B)** Counterclockwise rotation of the metaphase spindle in the π-EB1 H1299 cell on the left in response to an off-axis blue light pattern rotation by 1° every 40 s. Note that the spindle in the cell on the right is not exposed to blue light and does not directionally rotate during the observation time. Related to Fig. 3. Scale bar: 5 µm.

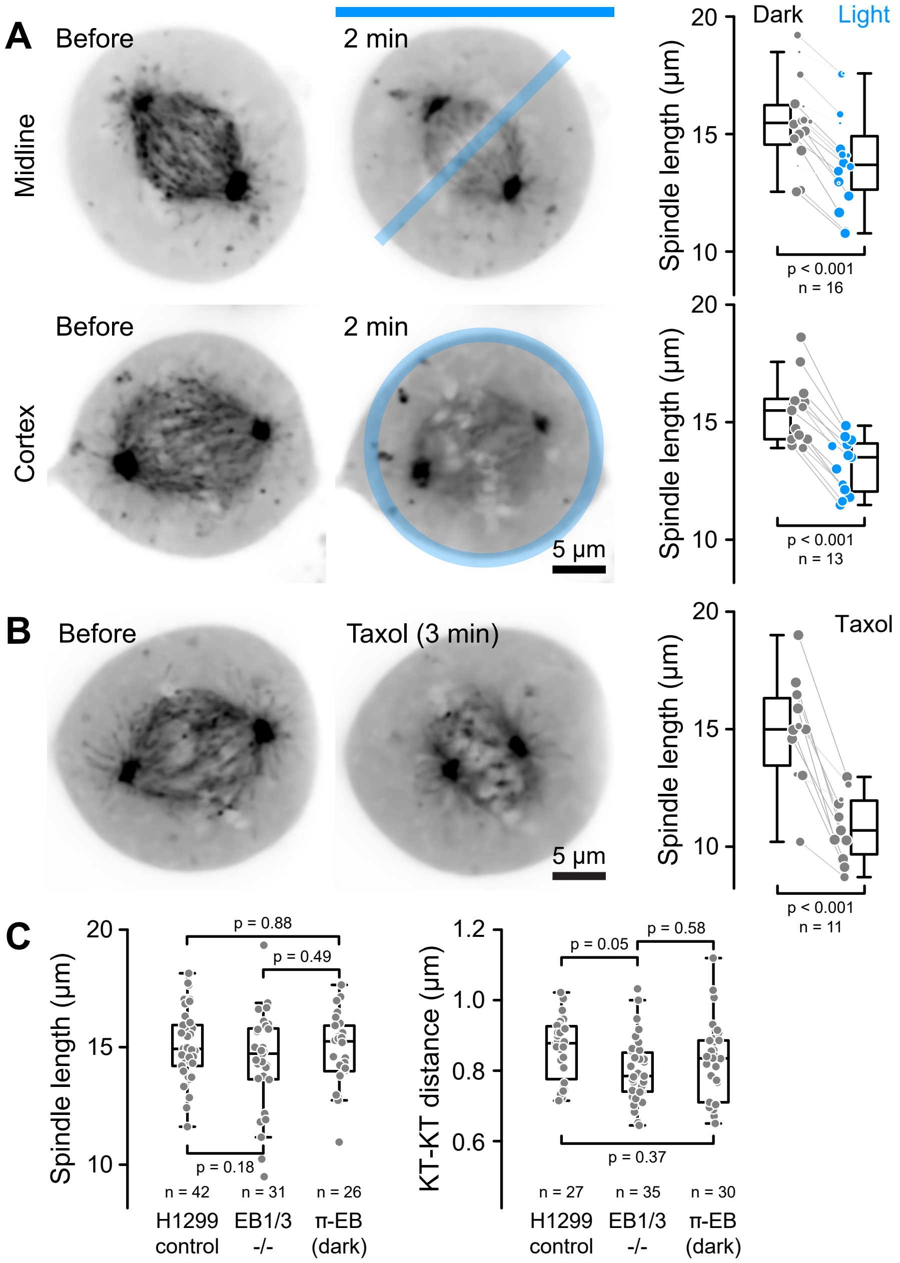


Dema et al., supplementary Figure 1

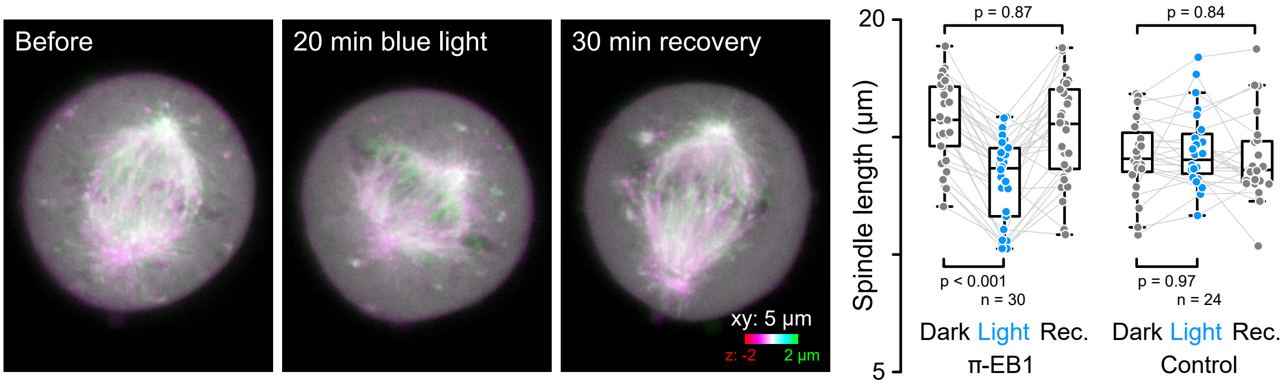


Dema et al., supplementary Figure 2

**A**

Neuronal (NP_004073 DCTN1 isoform 1) ...KVLKREGTDTTAKTSKLRGLKPKKAPTARKTTTRRPKPTRP...

Ubiquitous (NP_001128512 DCTN1 isoform 3) ...KVLKREGTDTTAKTSKL--------------------PTRP...

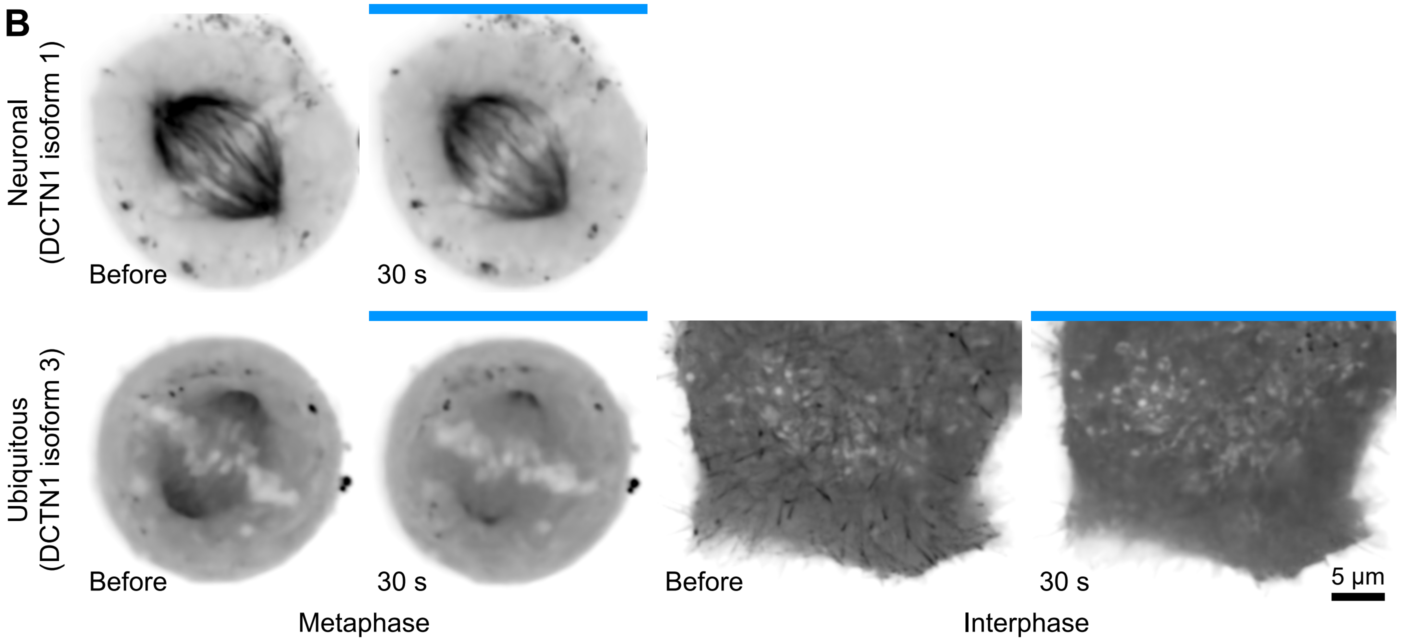


Dema et al., supplementary Figure 3
